# Ganglioside GM1 produces stable, short, and cytotoxic Aβ_40_ protofibrils

**DOI:** 10.1101/2022.12.28.522135

**Authors:** Manjeet Kumar, Magdalena I Ivanova, Ayyalusamy Ramamoorthy

## Abstract

Monosialoganglioside GM1-bound amyloid β-peptides have been found in patients’ brains exhibiting early pathological changes of Alzheimer’ s disease. Herein, we report the ability of non-micellar GM1 to modulate Aβ_40_ aggregation resulting in the formation of stable, short, rod-like, and cytotoxic Aβ_40_ protofibrils with the ability to potentiate both Aβ_40_ and Aβ_42_ aggregation.

---

Alzheimer’ s disease (AD) is associated with the gradual accumulation of cross-β amyloid fibrils of Aβ_40_ and Aβ_42_ in brains. Protein and peptide fibrillation is not due to the direct conversion of soluble monomeric form to insoluble amyloid fibrils. Instead, multiple intermediate processes may be involved, including the formation of prefibrillar or protofibrillar species that mainly act as primary or secondary nuclei for further fibril growth.^1^ These intermediate aggregates are considered to be the dominant cytotoxic forms of misfolded proteins/ peptides.^2–4^

Yanagisawa et al. identified a unique GM1 ganglioside-bound Aβ species in the brains of patients with early pathological changes associated with AD.^5,6^ Gangliosides, including GM1, are abundantly found in the neuronal membrane. Generally, free gangliosides are released in the extracellular region from the damaged cell membrane.^7,8^ Several studies have demonstrated that ganglioside micelle, or clusters in the membrane, catalyze toxic Aβ species’ formation.^9–17^ Additionally, free lipids have been shown to modulate the aggregation kinetics of Aβ peptides.^18–21^ This *in vitro* study is focused on unraveling the molecular processes underlying the early conformational changes of Aβ induced by the interaction with GM1, which have been linked to early AD onset. Our results show that Aβ_40_ produced stable, thioflavin T (ThT)-positive, cytotoxic, rod-like structures of diameter (22.5 ± 13.1 nm) and length (38 ± 20.3 nm) in the presence of GM1. Ganglioside induced Aβ_40_ species were able to catalyze the conversion of unfolded monomeric Aβ_40_ and Aβ_42_ to β-sheet rich structures.

The interaction of Aβ_40_ with non-micellar GM1 was monitored via ThT fluorescence assay (Fig.1A). The critical micelle concentration (CMC) of GM1 used in our study was calculated to be 79.4 ± 5.6 µM (Fig.S1). *In vitro* experiments carried out at physiologically relevant pH and temperature conditions, reported the formation of Aβ_40_ aggregates via a nucleation-dependent polymerization, typically displaying a sigmoidal curve with a prominent lag-phase, a growth phase, and an equilibrium phase.^22,23^ Our Aβ_40_ samples showed similar aggregation kinetics with a lag-phase (T_lag_) of around 8.8 hours (h). T_lag_ for the formation of β-sheet rich species in Aβ_40_ alone and GM1-containing samples were deduced from the ThT fluorescence data fitted to a sigmoidal function (Fig.1A).^24^ Interestingly, there was a significant reduction in the lag-phase of Aβ_40_ amyloid fibril formation and a slight increase in ThT fluorescence upon the addition of increasing concentration of GM1. A 1 µM GM1 was sufficient to reduce the lag-phase of 10 µM Aβ_40_ by 4.8 h. The decrease in the lag-phase and the increase in ThT fluorescence intensity suggested an acceleration of Aβ_40_ fibril formation. On the contrary, Chakravorty et al.,^18^ reported the inhibition of Aβ_40_ aggregation, which may be due to the use of frozen GM1 stock solution. To confirm the above observation, TEM images were acquired from 10 µM Aβ_40_ after incubation for 48 hours at physiological pH and temperature in the absence (Fig.1B) and the presence of 1 µM GM1 (Fig.1C, Fig.1D). Aβ_40_ alone showed the presence of typical long amyloid fibril of average diameter (10.6 ± 2.1), as previously reported.^25^ However, significantly shorter, rod-like fibrillar structures of varying width (average = 22.5 ± 13.1 nm) and length (average = 38 ± 20.3 nm) were observed in GM1 modulated Aβ_40_. Among the heterogeneous population of dominant short, rod-like GM1-modified Aβ_40_ species, a rare long fibril, most likely of non-GM1 bound Aβ_40,_ was also observed. The short, rod-like Aβ_40_ filaments bind more efficiently to ThT, indicating they have similar cross-β architecture as observed in most amyloid fibrils.^26,27^

Several studies have suggested ganglioside nanoclusters in neuronal membranes induce a conformational change in Aβ_40_.^28–31^ So, we were interested in examining if the Aβ_40_ can also undergo a secondary structure transition in the presence of GM1. Far-UV circular dichroism (CD) of 20 µM Aβ_40_ in the absence and presence of 2 µM GM1, monitored over 63 h at physiological pH and temperature, did not show any change except for the decrease in random-coil content, as evidenced by the reduction of negative ellipticity peak at 198 nm (Fig.2A, Fig.2B, Fig.2C). An immediate reduction in random-coil content in Aβ_40_ with no subsequent appearance of other secondary structure peaks as soon as GM1 was added suggests a fraction of Aβ_40_ rapidly forms insoluble aggregates/nuclei. This may explain the reduction in the lag-phase of GM1-containing Aβ_40_ in Figure 1. After that, we compared the molar ellipticity of Aβ_40_ with GM1 at 198 nm to the Aβ_40_ alone sample (Fig.2C, Fig.2D).

**Figure 1.**
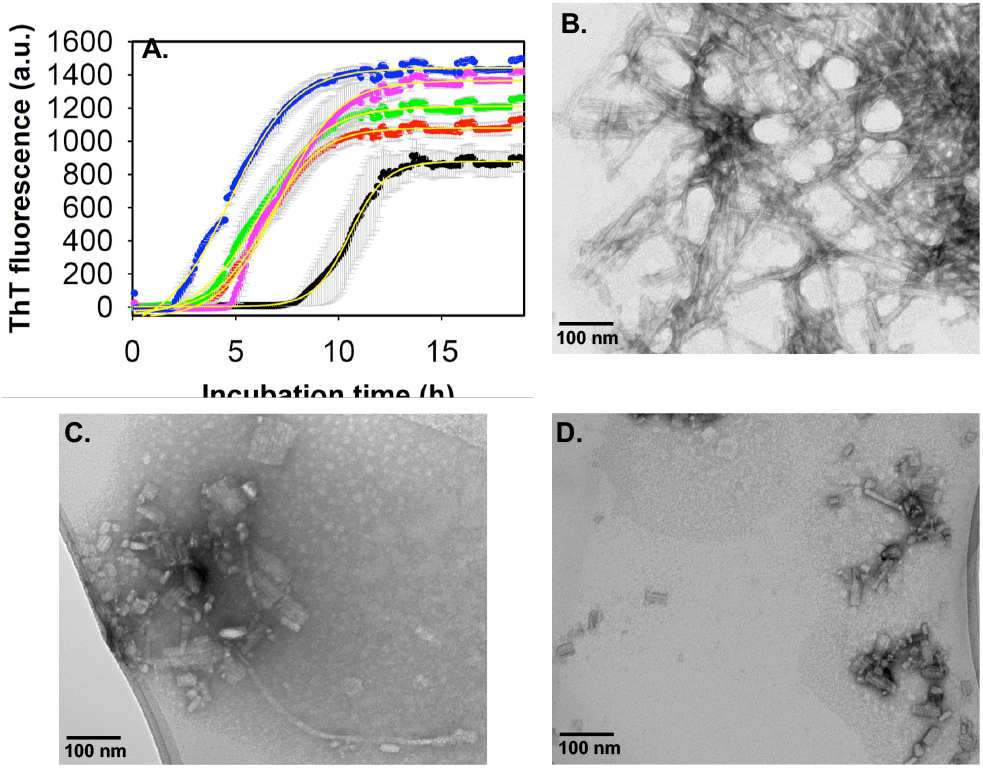
Probing the interaction between Aβ40 and GM1. (A) ThT fluorescence intensity of 10 µM Aβ40 alone (black) and with varying concentrations of GM1: 1 µM (red), 2.5 µM (green), 20 µM (magenta) and 40 µM (blue) in 20 mM phosphate buffer, pH 7.4 at 37° C, slow continuous shaking. The line of best fit (yellow, R2 =0.99) through the data points was obtained by global fitting the data with a sigmoidal function.24 TEM images of 10 µM Aβ40 alone (B), and in the presence of 1 µM GM1 (C, D), acquired after 48 h incubation at physiological pH and temperature conditions. (C) and (D) are the images of the same sample taken from two different positions in the TEM grid. Scale bars, as indicated. All other details are given in the supporting information.

**Figure 2.**
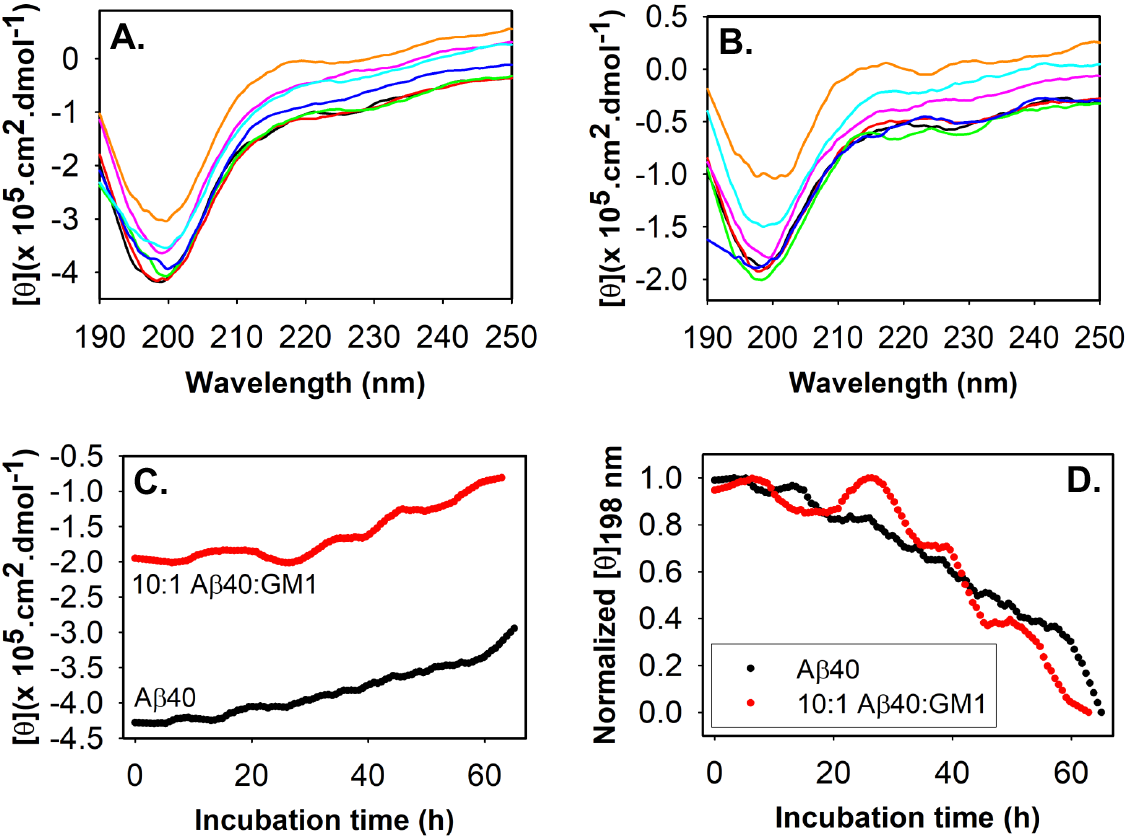
Comparing the rate of secondary structure change in Aβ40 during fibrillation in the absence and presence of GM1 via far-UV CD. Molar ellipticity of 20 µM Aβ40 (A) and 20 µM Aβ40 + 2 µM GM1 (B) measured over a period of about 63 h: black (0 h), red (10 h), green (20 h), blue (30 h), magenta (40 h), cyan (50 h) and mustard (65 h). Samples were measured in 20 mM phosphate buffer, pH 7.4 at 37° C. (C) Molar ellipticity of 20 µM Aβ40 (black) and 20 µM Aβ40 + 2 µM GM1 (red) at 198 nm over a period of^∼^63 h. (D) Normalized molar ellipticity at 198 nm of Aβ40 in the absence and presence of GM1 as indicated. All other details are provided in the supplementary information.

Both Aβ_40_ samples displayed a similar rate of change in the random coil during their aggregation. Overall, we infer that the nucleating species forming the cross-β sheet-rich structure are similar in both samples. Moreover, the apparent rate constant (k_app_) for the formation of cross-β sheet rich species in Aβ_40_ alone and GM1-containing samples were deduced from the ThT fluorescence data fitted to a sigmoidal function (Fig.1A).^24^ The k_app_ for 10 µM Aβ_40_ alone is 0.019/h, while the k_app_ for 10 µM Aβ_40_ in the presence of 1 µM, 2.5 µM, 20 µM, and 40 µM are 0.013/h, 0.011/h, 0.012/h and 0.01/h, respectively. The comparatively similar k_app_ for GM1-containing Aβ_40_ and the absence of long fibrillar species (Fig.1C) compared to Aβ_40_ alone suggested that GM1 inhibited the elongation of Aβ_40_ fibrils.

The ThT, TEM imaging, and far-UV CD results confirm that the short, rod-like Aβ_40_ species generated upon interaction with GM1, is an on-pathway stable intermediate, most likely protofibrils of fully matured fibrils of Aβ_40_. Interestingly, metastable protofibril of Aβ with the ability to alter the electrical activity of neurons, causing neuronal loss, has been reported earlier.^32–34^ We, therefore, performed a cell viability assay on differentiated human neuroblastoma SH-SY5Y cells treated with samples containing Aβ_40_ fibril, GM1-modulated Aβ_40_ protofibril, and GM1 alone (Fig.3). Sample containing 10 µM Aβ_40_ fibril alone did not show any significant cell cytotoxicity on differentiated SH-SY5Y cells, as reported earlier.^35^ However, we observed approximately 33% less cellular metabolic activity from differentiated SH-SY5Y cells upon incubation with GM1-induced Aβ_40_ protofibrils. A 1 µM GM1 (alone) incubated for 48 h showed no cell death. This observation implies that Aβ_40_ protofibrils are potentially cytotoxic.

**Figure 3.**
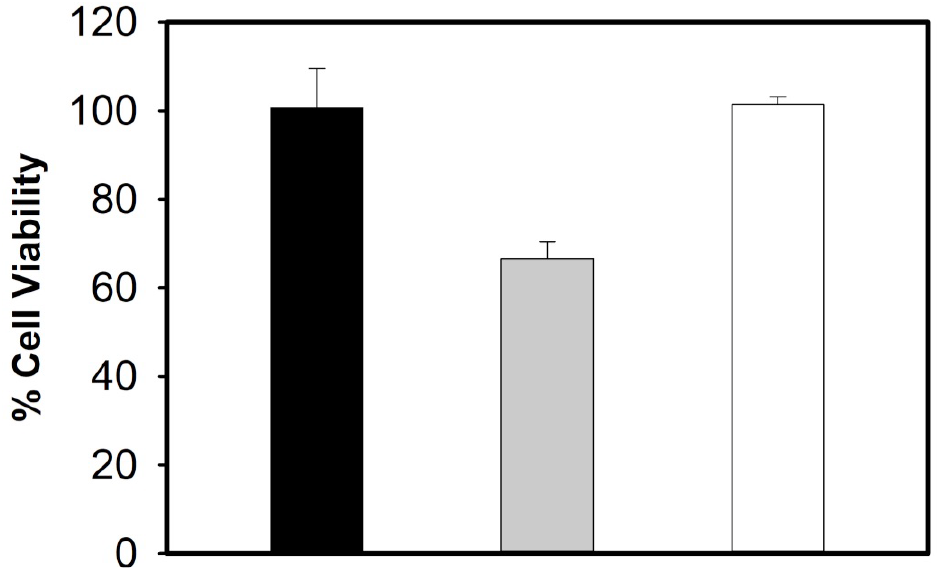
Evaluation of GM1-induced Aβ40 protofibril cytotoxicity. The MTT assay was used to measure differentiated human neuroblastoma SH-SY5Y cellular metabolic activity as an indicator of cell viability upon treatment with solutions containing 10 µM Aβ40 species without (black) and with (grey) 1 µM GM1 after 2 days of incubation. The white bar is of 2-day old 1 µM GM1 alone. Data is presented as the average ± standard error, calculated from 5 replicates (n = 5). Kruskal-Wallis test indicate that the result is significant at p < 0.05. See the supporting information for additional details.

Apart from the direct toxic effect displayed by the intermediates of many amyloid-forming proteins, as observed in this study, aggregating proteins can catalyze the conversion of other amyloid or neighboring proteins/peptides to aggregates^36^. The formation of amyloid fibrils from normal functional, misfolded, or unfolded proteins mostly follows nucleation-dependent polymerization similar to crystallization.^37,38^ The critical step in these processes is the generation of the smallest unit, often termed nuclei (seeds), that can promote the formation of amyloids by rapidly converting non-aggregated proteins/peptides into fibrils.^33^ There are two types of polymerization: homogenous polymerization, where both the nuclei and precursor proteins/peptides are the same, and heterogeneous polymerization (cross-seeded),^39^ in which the nuclei of one protein/peptide seed the aggregation of a different protein/peptide. Apart from primary nucleation, secondary nucleation, generally surface-catalyzed conversion of proteins/peptides to amyloid fibrils, can also lead to fibril proliferation.^40^ The rapid transformation of non-aggregated proteins/peptides through primary and secondary nucleation has been established as a central process explaining the “infectious” nature of prion proteins in prion diseases and the spread of pathogenic inclusions in many neurodegenerative disorders, including AD.^41–43^ Knowing the importance of nucleation in AD, we studied whether GM1-induced Aβ_40_ protofibril can catalyze the conversion of Aβ_40_ and Aβ_42_ (second major isomer generated from amyloid precursor protein involved in AD pathogenesis) to β-sheet rich structure. For seeding and cross-seeding experiments, we prepared the Aβ_40_ seeds by incubating 10 µM Aβ_40_ in the absence and presence of 1 µM GM1, without ThT dye in the amyloid-forming condition as described in the materials and method section (see the supplementary information). Using the ThT fluorescence assay, we studied the secondary nucleation process of Aβ_40_ by adding varying amounts of preformed Aβ_40_ fibril to 5 µM of monomeric Aβ_40_ under the quiescent condition at physiological pH and temperature (Fig.4A). Addition of 1%, 2.5% and 5% v/v preformed fibril as seed, significantly reduced the lag-phase of 5 µM Aβ_40_ aggregation kinetics. Meanwhile, additions of 1%, 2.5%, and 5% v/v preformed Aβ_40_ fibril to 5 µM of monomeric Aβ_42_ led to a spontaneous increase in ThT fluorescence, suggesting immediate Aβ_42_ fibril formation (Fig.4B). Seeding and cross-seeding of Aβ_40_ and Aβ_42_ respectively has earlier been observed with sonicated Aβ_40_ fibril.^44^ However, our study used unsonicated Aβ_40_ fibril as seeds. Similar to the secondary nucleation observed with Aβ_40_ fibril (Fig.4A, Fig.4B), the addition of preformed GM1-induced Aβ_40_ protofibril as seed (1%, 2.5% and 5% v/v) significantly reduced the lag-phase of 5 µM Aβ_40_ aggregation kinetics. Additionally, Aβ_40_ aggregates formed in the presence of GM1-induced Aβ_40_ protofibril bound more efficiently to ThT as indicated by higher ThT fluorescence intensity (Fig.4C). Furthermore, an immediate rise in ThT fluorescence was observed upon the addition of varying amounts of GM1-induced Aβ_40_ protofibril to 5 µM of monomeric Aβ_42_, which had a prominent lag-phase of ^∼^2 h without the seed (Fig.4D). That is, GM1-induced Aβ_40_ protofibril also catalyzed the aggregation of monomeric Aβ_40_ and Aβ_42_. As summarized in supplementary figure S2, we have shown the formation of short, rod-like intermediate of Aβ_40_, exceptionally stable protofibrils structure of average length of less than 40 nm upon interaction with ganglioside GM1. The GM1-induced Aβ_40_ protofibril reduced the viability of human neuroblastoma cells, SHSY5Y. Several studies, as mentioned earlier, and reviewed by Matsuzaki^9^, have shown that the interaction of GM1 in micellar or nanocluster forms with Aβ_40_ brings about an immediate α-helix/ β-sheet conformational change leading to the formation of toxic, long fibrillar amyloid-β species. However, our study found that non-micellar GM1 produces stable, toxic Aβ_40_ protofibrils of less than 40 nm size without undergoing α-helical transition upon interaction with the peptide. Potential implications of novel Aβ_40_ species reported in this study include: GM1-bound Aβ_40_ may be secreted efficiently due to its stability and smaller size into the extracellular space and act as a “seed” for amyloid-forming proteins/peptides, leading to a large-scale aggregation, apart from having a direct toxic effect to cells.^45,46^ Thus, our findings may explain the molecular basis for the early pathological changes observed in some patients with AD. In hindsight, the generation of stable, neurotoxic, protofibrillar assemblies of Aβ_40_ exhibiting prion-like properties to seed the aggregation of other proteins could contribute more to the neurodegeneration than the other observed oligomeric and amyloid fibrils. From a therapeutic perspective, blocking the formation of stable intermediates resembling those described here could successfully combat neurodegenerative diseases, including AD.

**Figure 4.**
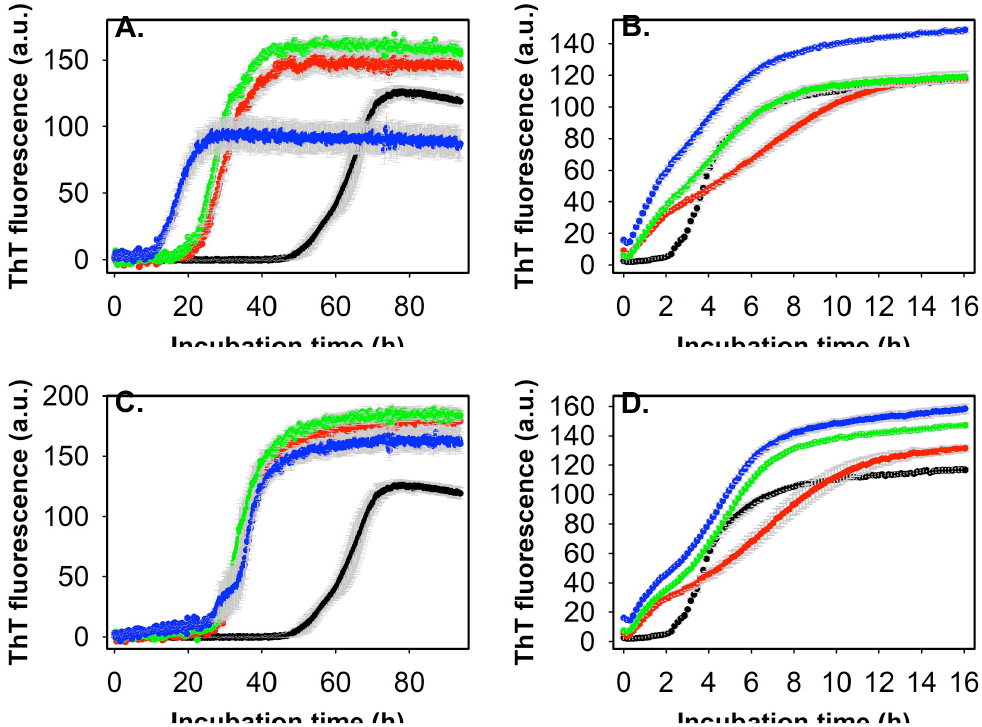
Seeding and cross-seeding of Aβ40 and Aβ42 with pre-formed Aβ40 aggregates. (A) ThT fluorescence profiles of 5 µM Aβ40 alone (black) and in the presence of 1% v/v (red), 2.5% v/v (green) and 5% v/v (blue) pre-formed Aβ40 amyloid fibril in 20 mM phosphate buffer, pH 7.4, 37° C under quiescent condition. (B) ThT fluorescence profiles of 5 µM Aβ42 alone (black) and in the presence of 1% v/v (red), 2.5% v/v (green) and 5% v/v (blue) pre-formed Aβ40 amyloid fibril in 20 mM phosphate buffer, pH 7.4, 37° C under quiescent condition. (C) ThT fluorescence profiles of 5 µM Aβ40 alone (black circle) and in the presence of 1% v/v (red circle), 2.5% v/v (green circle) and 5% v/v (blue circle) pre-formed GM1-induced Aβ40 protofibril in aforementioned conditions. (D) ThT fluorescence profiles of 5 µM Aβ42 alone (black circle) and in the presence of 1% v/v (red circle), 2.5% v/v (green circle) and 5% v/v (blue circle) pre-formed GM1-induced Aβ40 protofibril in above mentioned conditions. Appropriate blank was subtracted in each case. See the supporting information for additional details.

M.K. and A.R. designed the research, M.K. performed the experiments and processed the data, M.K., M.I. and A.R. analyzed the data, M.K., M.I. and A.R. wrote the manuscript, and A.R. directed the project.

## Acknowledgment

This study was supported by the National Institutes of Health Grants (AG048934 and DK13221401 to A.R.).

