## Supporting Information for "Ganglioside GM1 produces stable, short, and cytotoxic Aβ_40_ protofibrils"

1. **Determination of critical micelle concentration (CMC) of GM1 ganglioside using the pyrene fluorescence assay.** The reported CMC value of GM1 in aqueous solution ranges from ~10 nM[1], ~20 nM[2], < 1 µM[3], ~28 µM[4], ~85 µM[5] to ~100 µM[6]. The large variations in the reported CMC of GM1 may be due to different sample preparation methods and techniques used to measure the CMC value. To determine whether the GM1 used in our current study is in a micellar or non-micellar form, we measured the CMC value through the pyrene fluorescence assay as described by Aguiar et al.[7] and Chakravorty et al.[6] The CMC value of GM1 in our condition is estimated to be 79.4 ± 5.6 µM as shown in figure S1.

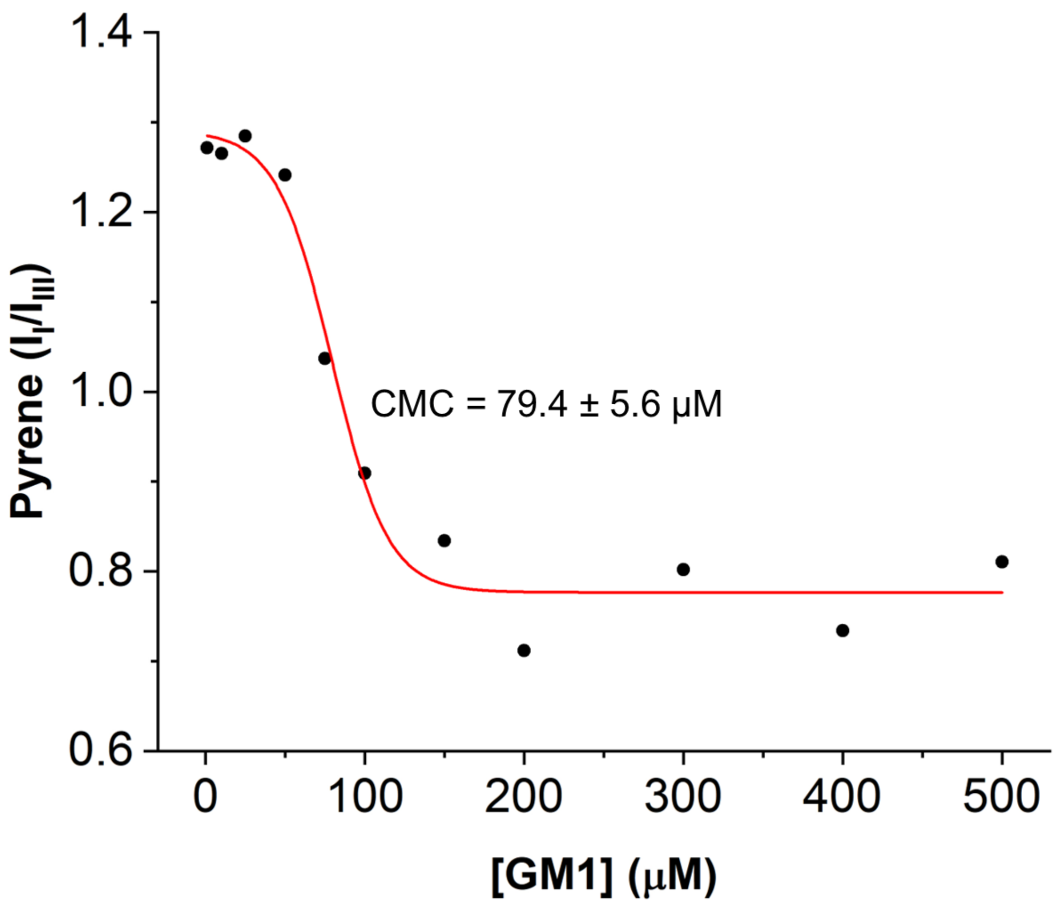


**Figure S1. Pyrene fluorescence assay for measuring the CMC of GM1.** The pyrene fluorescence intensity ratio of peak I (373 nm) and peak III (384 nm) (i.e., $I_I/I_{III}$) was plotted against increasing concentrations of GM1 (from 1 to 500 µM). CMC of GM1 was calculated as the midpoint of the Boltzmann-type sigmoid fit (red line, $R^2$ = 0.97), implemented within the OriginLab 2022 software package, to the above data.

### 2. Summary of the effect of GM1 on Aβ-40 amyloid fibril formation.

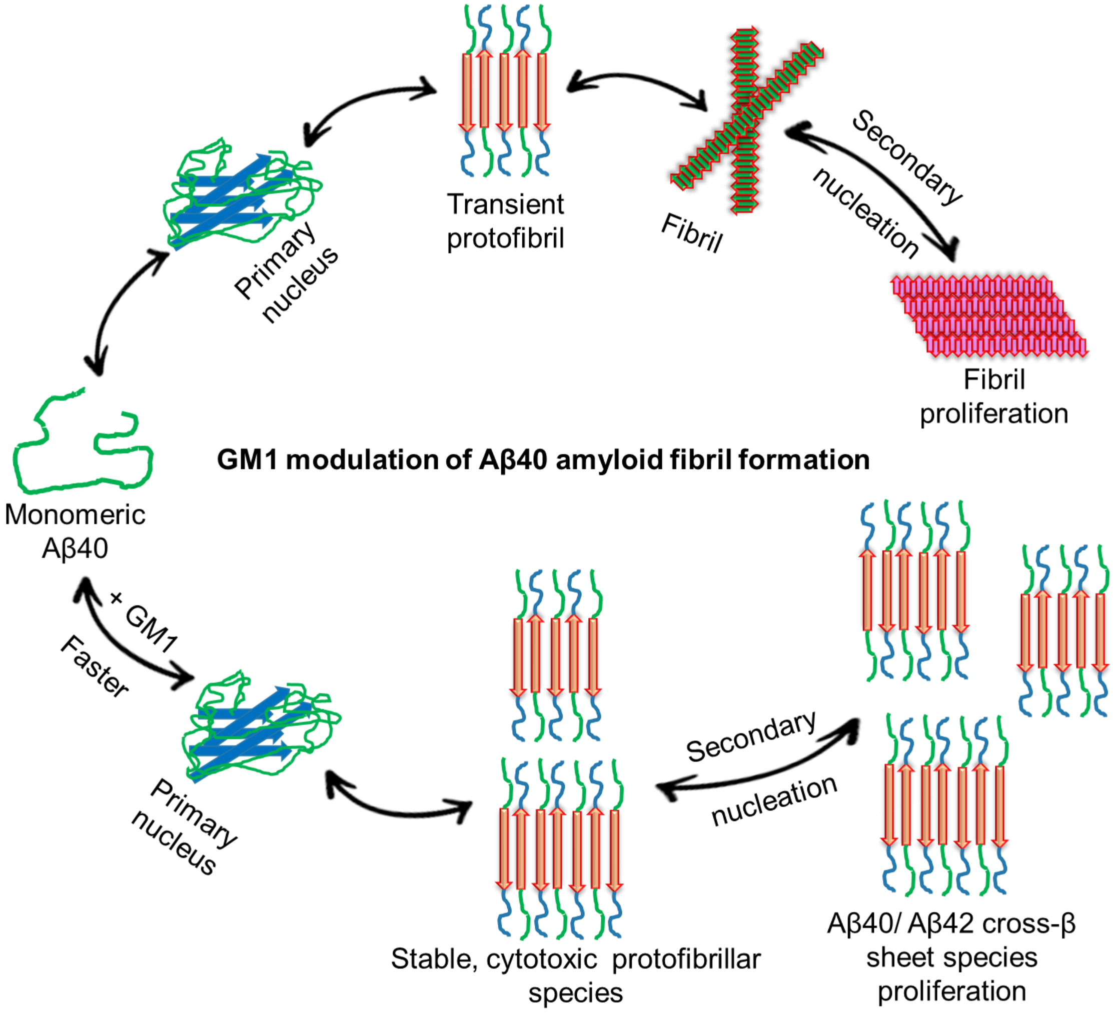


**Figure S2. Schematic representation of the effect of GM1 on $A\beta_{40}$ amyloid fibril formation.** At physiological pH, temperature, and under constant agitation conditions, $A\beta_{40}$ slowly generates amyloidogenic precursor (primary nuclei), leading to the formation of transient protofibrillar species, which subsequently form amyloid fibrils. The $A\beta_{40}$ fibril surface can catalyze the conversion of monomeric $A\beta_{40}$ and $A\beta_{42}$ to aggregates leading to fibril proliferation. However, in the presence of GM1, $A\beta_{40}$ amyloidogenic precursor formation is faster, leading to the formation of stable, protofibrillar species. The GM1-modulated $A\beta_{40}$ protofibril is neurotoxic with the tendency to proliferate $A\beta_{40}$ and $A\beta_{42}$ cross-β sheet rich aggregates formation.

### Materials and Experimental Procedures

#### *Materials*

Amyloid-β peptides ($A\beta_{40}$ and $A\beta_{42}$) were purchased from Vivitide (Gardner, MA). GM1 ganglioside (Ovine Brain, Lot: 860065P-5MG-H-021) was obtained from Avanti Polar Lipids (Alabaster, AL). Unless otherwise stated, all other chemicals were purchased from Sigma Aldrich (St. Louis, MO).

#### *GM1, Pyrene, Aβ Peptide, and Thioflavin T (ThT) Stock Preparation*

In a 1.5 mL Eppendorf tube, 1 mg of GM1 lipid was mixed in 100 µL methanol. A solvent such as methanol has recently been commonly used to solubilize gangliosides, including GM1.[8,9] Once fully dissolved, the lipid solution was aliquoted and dried under a stream of dry nitrogen gas. Residual solvents were removed in a vacuum chamber for at least 1 hour and stored at -20 °C for further use. Immediately before use, dried lipid was rehydrated in 20 mM sodium phosphate buffer, pH 7.4, and mixed by vortexing to give a final concentration of 64 µM. 0.98 mM stock solution of pyrene was prepared by dissolving 1 mg of pyrene crystals in 5 mL of 100% ethanol. Powder samples of Aβ peptides were dissolved at 1 mg/mL in hexafluoroisopropanol (HFIP) and incubated for 1 hour to monomerize any existing aggregates. The peptides were aliquoted, lyophilized overnight, and stored as a powder sample at -20 °C. Immediately before the use in further experiments, Aβ powders were dissolved in 20 mM sodium phosphate buffer, pH 7.4, to make ~200 µM solution. Exact concentrations were confirmed by UV absorbance on a nanodrop (DeNOVIX, DS-11$^{+}$ Spectrophotometer), using the molecular weights of $A\beta_{40}$ and $A\beta_{42}$ as 4.3 kDa and 4.5 kDa, respectively, and 1490 $M^{-1}$ $cm^{-1}$ as the molar extinction coefficient at 280 nm. 1 mg ThT was dissolved in 1 mL 20 mM sodium phosphate buffer, pH 7.4 and the exact concentration was estimated by measuring the absorbance at 412 nm on a transparent 96-well plate (Fischer Scientific) in a Biotek Synergy 2 plate reader and using the molecular weight of ThT as 318.86 g and the molar extinction coefficient as 36,000 $M^{-1}$ $cm^{-1}$.

*Determination of the CMC of GM1 via pyrene fluorescence assay*

A fresh 6.4 mM stock solution of GM1 was prepared in 20 mM sodium phosphate buffer, pH 7.4, using the method described above. Different concentrations of freshly prepared GM1 ranging from 1 µM to 500 µM were mixed with 1.5 µM of pyrene in semi-micro disposable fluorescence cuvettes (fireflysci, type-9FLP, 10 mm pathlength). Pyrene fluorescence emission spectra at different concentrations of GM1 were recorded from 350 nm to 450 nm (emission slit-width = 5 nm) after excitation at 334 nm (excitation slit-width = 1 nm) in FluoroMax4® from Horiba Scientific® at 37 °C. The CMC of GM1 was calculated from the pyrene fluorescence intensity ratios obtained from peaks I and III (I/III) at 373 and 384 nm, respectively, as described by Aguiar et al.[7] and Chakravorty et al.[6]

*Determining the effect of GM1 on $A\beta_{40}$ amyloid fibril formation*

10 µM $A\beta_{40}$ in 20 mM sodium phosphate buffer, pH 7.4, 37 °C, was incubated in the absence and presence of varying concentrations of GM1 in 384-well microplates (Greiner bio-one black, clear bottom, non-binding, low volume, Lot: E22053E7), with continuous, slow shaking (frequency = 17 Hz, i.e., 1020 cycles/ min). The fibrillar aggregation of $A\beta_{40}$ was monitored via ThT fluorescence assay using specific filters of the Biotek Synergy 2 microplate reader. A transparent sealing film was used to prevent solvent evaporation. ThT fluorescence emission was measured using a 440/40 nm filter for excitation and a 485/20 nm filter for emission. A 1:0.33 peptide: ThT molar ratio was used for all the amyloid fibril kinetic assays.[10] The fluorescence spectra of $A\beta_{40}$ samples were plotted by subtracting the appropriate blanks (that exhibited negligible ThT fluorescence) as the average value of 3 replicates. The blank samples had varying concentrations of GM1 and ThT but no peptide. Standard error was calculated using the following formula.

$$Standard\ error = Standard\ deviation\ of\ the \frac{samples}{\sqrt{number} of\ replicates} \quad [1]$$

All aggregation curves were fitted to a sigmoidal function (global fitting, $R^2 = 0.99$) as described by Morris et al.,[11] implemented within the SigmaPlot 12.0 software package to determine the lag-phase ($T_{lag}$) and the apparent rate constant ($k_{app}$).

*Transmission electron microscopy*

5 µL of samples (prepared as described above) were transferred to ultrathin holey carbon support film, copper 400 mesh grids, for 2 min. The samples were blot-dried using Whatman filter paper, washed 3 times with Milli-Q water, and negatively stained with uranyl acetate (2% w/v; Electron Microscopy Sciences) for another 2 min followed by a quick wash with milli-Q. Samples were dried overnight and examined using a JEOL JEM 1400 plus transmission electron microscope at a 80 kV accelerating voltage under a magnification of 25,000 x. Particle sizing was performed using the ImageJ software (NIH). The diameter and length of the $A\beta_{40}$ aggregates are reported as mean ± standard deviation (n > 15).

*Real-time far-UV circular dichroism spectroscopy*

The real-time far-UV CD spectra of 20 µM $A\beta_{40}$ in the absence and presence of 2 µM GM1 in 20 mM phosphate buffer, pH 7.4, 37 °C was collected over a period of ~ 63 to 65 h in a 0.01 cm path length cuvette (QS high-precision cell, Hellma Analytics) sealed with a sealing tape in a JASCO J-1500 CD spectrometer equipped with a Julabo AWC100 water bath following a similar method as described Kumar et al.,[12] The approximate scan time for each spectrum was 4.016 min, which was collected over a wavelength range of 180-260 nm with a data pitch of 0.2 nm, a data integration time of 2 s, a bandwidth of 1.5 nm, a delay time of 0 min, waiting time of 0 min, and a scan rate of 20 nm/min. 2 µM GM1 (alone) in 20 mM phosphate buffer, pH 7.4, 37 °C showed no noticeable spectrum. The molar ellipticity of $A\beta_{40}$ samples was smoothed using the smoothers function implemented within the SigmaPlot 12.0 software package.

*Cell Viability Assay*

100 µL of 60,000 SH-SY5Y cells/mL (ATCC, catalog # CRL-2266, Lot number: 63724189) were plated in a 96-well plate and differentiated in Neurobasal-A supplemented with B27, GlutaMAX, 1% penicillin/streptomycin, and 10 µM retinoic acid in a humidified incubator at 37 °C with 5% $CO_2$ for 10 days. The used media was suctioned out and replaced with fresh 100 µL differentiation media on every alternate day. On the 11$^{th}$ day, 50 µL of fresh differentiation media mixed with 50 µL of the samples containing preformed 20 µM $A\beta_{40}$ aggregates formed in the absence and presence of 2 µM of GM1, and 2-day old 2 µM of GM1 (alone) were added to the cells and incubated for 48 hours. 20 mM sodium phosphate buffer, pH 7.4, was used as the negative control, and 2% SDS was used as a positive control for cell assay. On day-12, MTT (3-(4,5-dimethylthiazol-2-yl)-2,5-diphenyl-2H-tetrazolium bromide) cell proliferation assay (Promega, G4001) was performed to determine the toxicity of the aforementioned $A\beta_{40}$ samples following the manufacturer's protocol. 15 µL of the MTT dye solution was added to each well and incubated at 37 °C for 2 hours in a humidified $CO_2$ incubator, followed by 100 µL solubilization solution. The absorbance of each well was then measured both at 570 nm and 650 nm (background). The average cellular viability value of the negative control (20 mM phosphate buffer, pH 7.4) after background subtraction was taken as 100%. Values reported are the average of five independent trials, and the standard error was calculated as discussed above in equation [1]. MTT assay was performed twice for $A\beta_{40}$ aggregates (samples prepared from 2 batches of $A\beta_{40}$ peptides from the same supplier) and GM1 (same batch).

*Amyloid seeding/ cross-seeding experiments*

For seeding experiments, 5 µM of fresh $A\beta_{40}$ in 20 mM phosphate buffer, pH 7.4, was incubated in a Biotek Synergy 2 plate reader (no shaking) at 37 °C, either without (unseeded) or with (seeded, 1% v/v, 2.5% v/v, and 5% v/v) $A\beta_{40}$ aggregates formed in the absence and presence of GM1. The aggregation was performed in a 384-well microplate. For cross-seeding experiments, 5 µM of fresh $A\beta_{42}$ in 20 mM phosphate buffer, pH 7.4, was incubated at 37 °C either without (unseeded) or with (seeded, 1% v/v, 2.5% v/v, and 5% v/v) $A\beta_{40}$ aggregates formed in the absence and presence of GM1.

$A\beta_{40}$ aggregates (seeds) were formed by incubating 10 µM peptide in the above-mentioned amyloid-forming conditions, without ThT dye for 48 hours in the absence and presence of 1 µM GM1. Control $A\beta_{40}$ samples in the absence and presence of GM1 and ThT were also set up in the same microplate to track the aggregation. The *in situ* aggregation was monitored via ThT fluorescence, as described above. Briefly, 1.7 µM for (unseeded), 1.8 µM (1% v/v seed), 1.95 µM (2.5% v/v seed), and 2.2 µM (5% v/v seed) of ThT was added to the above samples. ThT fluorescence of 1.7 µM for (unseeded), 1.8 µM (1% v/v seed), 1.95 µM (2.5% v/v seed), and 2.2 µM (5% v/v seed) ThT without the fresh monomeric peptides were considered as blank. The fluorescence spectra were plotted by subtracting the appropriate blanks as the average value of 3 replicates. Standard error was calculated as mentioned above.

**Keywords:**

Alzheimer's disease, Amyloid-β, Ganglioside (GM1), Protofibril, Seeding, Cross-seeding

**Abbreviations**

| | |
|---|---|
| AD | Alzheimer's Disease |
| Aβ | Amyloid-beta |
| GM | Ganglioside |
| ThT | Thioflavin T |
| $T_{lag}$ | Lag-phase |
| $k_{app}$ | Apparent rate constant |
| TEM | Transmission Electron Microscopy |
| CD | Circular Dichroism |
| MTT | (3-(4,5-dimethylthiazol-2-yl)-2,5-diphenyl-2H-tetrazolium bromide) |

12 M. Kumar, S. L. Teakel, C. Swarbrick, I. S. Chowdhury, D. C. Thorn, M. Sunde, J. A. Carver, J. K. Forwood, C. J. K Forwood and J. A. Carver, *FEBS J*, , DOI:10.1111/FEBS.16795.
